# Characterization of transcriptomic changes in the neurovascular unit of Alzheimiers transgenic mouse models using digital spatial profiling

**DOI:** 10.1101/2025.03.03.640886

**Authors:** Vrishali S. Salian, Kevin J. Thompson, Xiaojia Tang, Val J. Lowe, Krishna R. Kalari, Karunya K. Kandimalla

## Abstract

Alzheimer’s disease (AD) affects 40 million individuals globally and is characterized by the accumulation of amyloid-beta (Aβ) proteins, which aggregate and form plaques.

BBB dysfunction drives AD cerebrovascular pathology and BBB integrity is maintained by neurovascular unit (NVU). Specifically, within the NVU, the cerebral endothelial cells maintain vascular homeostasis. In this study, we isolated endothelial-enriched regions of interest (ROIs) using the Nanostring GeoMx digital spatial profiler and employed a deconvolution model to evaluate transcriptomic changes. We observed dysregulation of cellular signaling potentially disrupting the APP+ BBB integrity. Analysis of ligand-receptor pairings that are the foundation of the NVU intercellular signaling indicated that the endothelial vasculature completes a feedback loop with the NVU to the regulating astroctyes. Further, we identified potentially antagonistic signaling roles for opioid receptor species that should be further investigated for potential therapeutic targets.

**Graphical Abstract:** 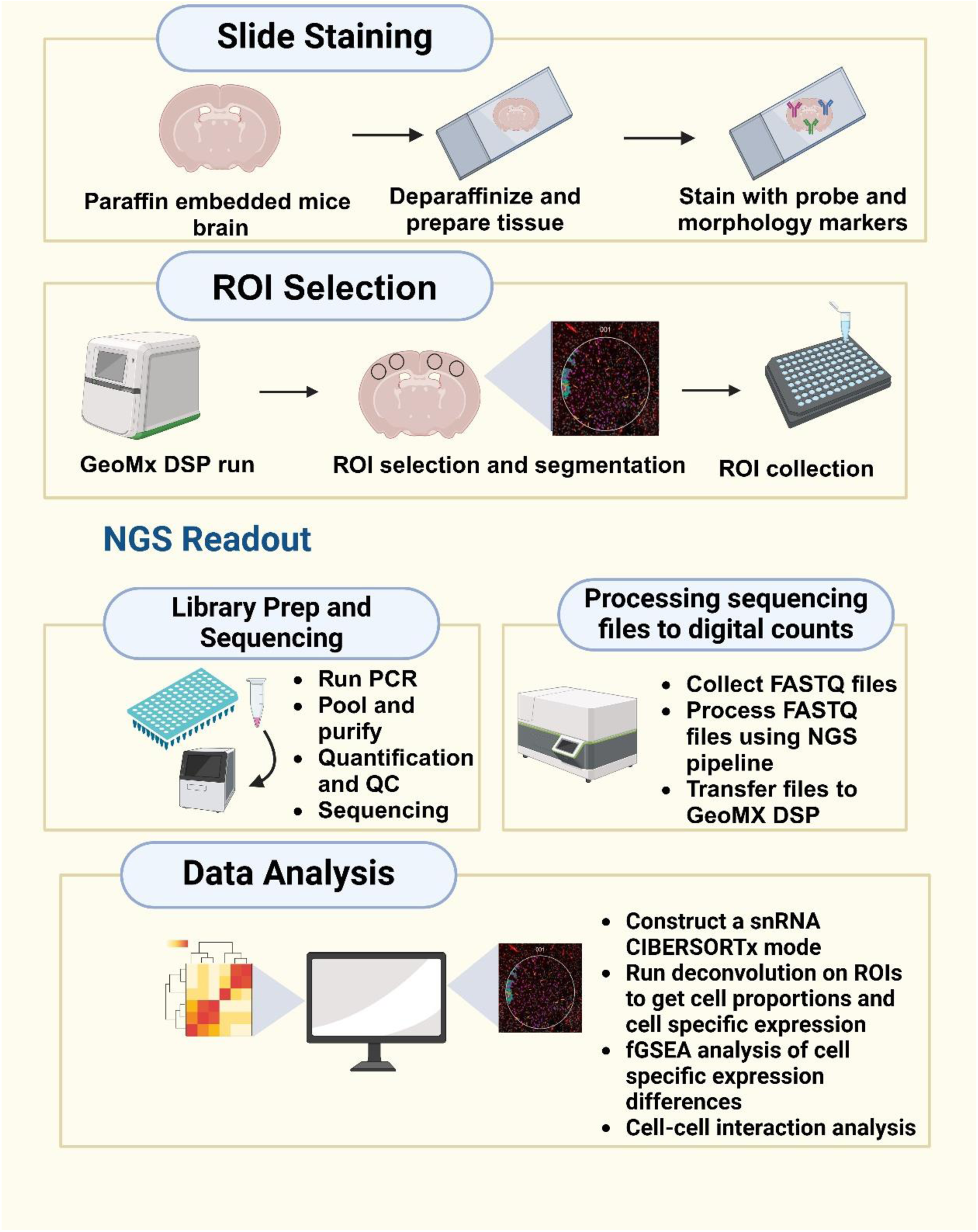

## Introduction

Alzheimer’s Disease (AD) is characterized by slow and progressive neurodegeneration and is reported to account for 60-70% of clinical dementia cases. AD currently affects an estimated 40 million individuals worldwide, with projections indicating that this number may double by 2030 and more than triple by 2050[1]. The pathological hallmarks of AD include the aggregation of soluble amyloid-beta (Aβ) to amyloid plaques and its subsequent accumulation in the brain parenchyma, in addition to the hyperphosphorylation of tau protein[2]. Moreover, cerebrovascular pathology plays a critical role in the progression of AD, as it disrupts the integrity of the blood-brain barrier (BBB) and is emerging as a pathological hallmark of AD that can be detected before the onset of cognitive impairment. The BBB serves to isolate the brain from systemic circulation, which makes the human brain vasculature a crucial point of clinical investigation. The exposure of Aβ to the BBB precipitates cerebrovascular pathology and constitutes one of the early pathophysiological changes associated with AD.

The integrity of the BBB is upheld by a functional neurovascular unit (NVU), which comprises vascular endothelial cells, pericytes, astrocytes, and neurons.

Collectively, these cell types protect the brain from systemic toxins and agents, maintain the homeostatic environment, and fulfill the metabolic demands of the cerebrovasculature, thereby facilitating proper neuronal function[3]. Among these cell types, endothelial cells, which constitute the inner lining of the BBB, are essential for maintaining vascular homeostasis. However, in the presence of Aβ, these cells become increasingly dysfunctional, contributing to endothelial inflammation and impaired clearance of toxic metabolites, including Aβ[4]. There is a rising interest in the maintenance of the BBB as it relates to both, the causative associations with amyloid and tau accumulation, and the therapeutic delivery of treatment agents. Despite the growing recognition of endothelial dysfunction in the pathology of AD, the molecular mechanisms underlying this dysfunction remain poorly understood. Studies investigating the mechanisms that contribute to BBB disruption have primarily focused on bulk-tissue analyses, which are likely to obscure the complex interactions among endothelial cells and other cell types within the NVU[5]. Advances in single-cell and single-nucleus RNA sequencing have enabled researchers to obtain comprehensive transcriptional data. Nevertheless, challenges remain in isolating the cells that comprise the BBB’s NVU using single-cell RNA sequencing, often resulting in the loss of RNA and/or DNA molecules during physical separation[6].

The Nanostring GeoMx® platform provides researchers with the capability to assay the whole transcriptome of rare and difficult-to-isolate cell types and their surrounding tissues with fewer processing steps designed to preserve RNA integrity. In this manuscript, we employed the GeoMx platform to assess transcriptional differences in the BBB vasculature between APP+ mouse models and their wild-type counterparts. We isolated endothelial-enriched regions of interest (ROIs) and determined that, although endothelial enrichment was achieved, additional NVU cell types contributed to the observed differences. To further elucidate the endothelial differences underlying the APP+ genotype, we developed a deconvolution model based on single-cell data from similarly aged mice. The sample-specific deconvolution of the collected ROIs was evaluated for differential expression, and we further investigated the cellular communications that underlie the NVU.

### Methods Animals

Eleven-month-old APP_swe_/PS1_deE9_ transgenic mice (APP+) (MMRRC Strain # 034832-JAX-HET), their wild-type (WT) littermates (APP-) (MMRRC Strain # 034832-JAX-WT), and B6SJLF1/J (MMRRC Strain # 100012), were purchased from the Jackson Laboratory (Bar Harbor, ME). A sample size of three male and female mice (n=3) was used for each group. All twelve mice were housed in the Mayo Clinic animal care facility under standard conditions with access to food and water *ad libitum.* The studies were conducted in accordance with the National Institutes of Health guidelines for the care and use of laboratory animals and protocols approved by the Mayo Clinic Institutional Animal Care and Use Committee (Mayo IACUC #A00006176-21).

### Sample preparation

The mice were transcardially perfused with cold phosphate-buffered saline (PBS) to flush out blood from the vasculature. Subsequently the mice were treated with a perfusion of 4% paraformaldehyde (PFA) in PBS to fix brain tissues. After perfusion, the brains were harvested and post-fixed in 4% PFA at 4°C for 24 hours. Following fixation, the brains were dehydrated through a graded ethanol series, cleared in xylene, and embedded in paraffin for histological processing and further analysis. Samples were processed in accordance with Nanostring’s (Seattle, Washington, USA) manual slide preparation guide. 5 μm thick sections of formalin-fixed, paraffin-embedded (FFPE) brain tissues were baked at 60°C for 2 hours. Following which they were deparaffinized with CitriSolv (Decon Laboratories (King of Prussia, PA, USA),1601H) rehydrated with 2X 100% ethanol and 1X 95% ethanol, then washed in 1X PBS. Target antigen retrieval was performed by placing slides in a staining jar containing tris ethylenediamine tetra acetic acid (EDTA, pH 9) and incubated at low pressure at 99 °C for 20 minutes, followed by a 5 min wash in PBS. The slides were then placed in a staining jar with 0.1 µg/mL proteinase K (ThermoFisher Scientific (Waltham, Massachusetts, USA), AM2548) and incubated at 37 °C for 10 minutes. Subsequently, the slides were post-fixed in 10% neutral buffered formalin (NBF) at room temperature, following which they were washed 2X in NBF stop buffer (StatLab (McKinney, TX, USA), 28600-1) and one wash with 1X PBS.

### Hybridization with RNA detection probes

The brain sections were incubated overnight at 37°C with Mouse Whole Transcriptome Atlas RNA detection probes diluted in buffer R (NanoString Technologies) using a RapidFISH hybridization oven (Boekel Scientific). The oligonucleotide probes contained a sequence complementary to the target mRNA, connected to a sequencing oligonucleotide via a ultraviolet photocleavable linker. HybriSlip Hybridization Covers (Grace Bio-Labs, Inc (Bend, OR, USA) 714022) were used to cover the slides during incubation. Slides were washed twice with a stringent wash buffer containing Saline-Sodium Citrate (SSC) buffer and formamide at 37 °C and then twice with SSC buffer following which, the slides are placed in a humidity chamber and blocked with buffer W (NanoString Technologies) for 60 minutes. To visualize endothelial cells during the GeoMx Digital Spatial Profiler scan, the slides were incubated overnight with CD31 primary antibody at 1:50 dilution. Subsequently, the anti-rat AF647 secondary antibody was applied and slides incubated for 60 minutes at room temperature. Slides were washed 2X in 2XSSC and stained with SYTO 13 green, fluorescent nucleic acid stain (5 µM, ThermoFisher, Cat# S7575). The slides were placed on the Nanostring GeoMx Digital Spatial Profiler (DSP) instrument and scanned at 20X magnification to fluorescently visualize the CD31 staining and cell nuclei.

### Imaging

Area of illumination (AOI) masks were created using the GeoMx DSP Control Center software by utilizing the fluorescence intensity of the CD31 staining in the cortex and hippocampus. The AOIs were manually selected using circular or rectangular regions measuring around 660 µm in diameter or 660 µm x 700 µm, respectively. The thresholding of each segment was adjusted for each AOI to accommodate staining heterogeneity, and segments were selected for CD31 positive regions. The segment area and number of nuclei in each AOI was determined using the GeoMx software, which counts cells in the Syto 13 channel, with a sampling of 100 cells were collected per AOI. The DSP instrument exposed each AOI with UV light permitting the photocleavage of a 66 bp-long DNA barcode from each oligonucleotide probe that was hybridized to an mRNA target. Following UV exposure, the indexed oligonucleotides from the entire AOI were collected and deposited in a 96-well plate. This process of localized UV exposure and UV cleaved-oligo collection was repeated for each AOI. The 96-well plate was sealed with a permeable membrane and left on the bench overnight to allow for complete sample evaporation. Subsequently, the samples were resuspended with 10uL of DEPC water and incubated at room temperature for 10 minutes.

### Library preparation

The construction of the sequencing libraries was completed in accordance with the manufacturer’s instructions (GeoMx—NGS Readout Library Prep User Manual, MAN-10153-03 FEB 2023) to include specific Illumina i5/i7 dual indexing primers. To each PCR reaction 2 µL of the GeoMx Seq Code PCR Master Mix, 4 µL of the GeoMx SeqCode, and 4 µL of the rehydrated samples was added. Thermocycling conditions for the PCR run were 37°C for 30 min, 50°C for 10 min, 95°C for 3 min; 18 cycles of 95°C for 15 s, 65°C for 1 min, 68°C for 30 s; and 68°C for 5 min. The PCR reactions were pooled (4 μL of each individual PCR product) and purified using two rounds of AMPure XP (Beckman Coulter, Cat#A63880) bead cleanup, in accordance with the manufacturer’s instructions. Following which, the samples were resuspended in elution buffer (Tris-HCl 10 mM with 0.05% Tween-20, pH 8.0).

### Library quantification and quality assessment

The library pools for the collection plates were quantified using the Qubit dsDNA HS kit assay (Invitrogen, cat. Q32854) on a Qubit 3.0 Fluorometer (Invitrogen). A 2200 TapeStation System (Agilent) High Sensitivity D1000 ScreenTape Assay (Agilent, cat. no. 5067-5584, 5067-5585) was performed to assess the quality of the library and to confirm the amplicon size, which is expected to be 162 bp.

### Sequencing

The sequencing depth was determined based on the manufacturer’s recommendation of 100 reads per square micron of Area of Illumination (AOI). Elution Buffer (Tris-HCl 10 mM with 0.05% Tween-20, pH 8.0; Teknova, cat. no. T1485) was used to normalize the library pool to 1 nM. Using an Illumina NextSeq 2000, 100 cycles, paired-end sequencing was carried out according to the following parameters: paired-end with reads 2 x 27 bp; 5% PhiX spike-in; dual-indexing with Index 1 (i7) 8 bp and Index 2 (i5) 8 bp. Upon sequencing completion, BCL files were converted to FASTQ files, which were uploaded to the GeoMx DSP instrument.

### ROI Analysis

Regions of interest (ROIs) were extracted from 12 mice brains for subsequent analysis. The 133 ROIs represented a combination of APP+/APP-and male/female mice. The evaluation of ROIs was conducted with one exception, whereby the limit of quantification (LOQ) LOQ geometric standard deviations was set to 1.5. Differential expression analyses were performed using the limma package (version 3.58.1)[7], with adjustments made for gender differences among the mice. UCell analysis[8] was conducted to evaluate whether ROI expression profile originated from a harvested endothelial source.

### Application of Deconvolution Methods

The single cell transcriptional data from 8 aging mice (21-22 months) was obtained from <u>GSE129788</u> [9]. The transcriptional data for 20,688 cells from 14 cell clusters, herein reported as cell types (required > 100 single cell samples) was obtained from the 8 C57BL/6J mouse brains representing 14,699 genes. Single cell data was evaluated with UCell for 5 existing NVU cell types of signatures (astrocyte, neurons, endothelial, microglia, and oligiodendrocytes) [10].

We then constructed a single-cell gene expression matrix as the references data set for deconvolution with CIBERSORTx [13]. Among the 14 cell types, under-represented cell types were removed, and the rest were consolidated into 6 cell types for a balanced ratio between sample size and number of targeted cell types. Cell type specific gene features were selected by evaluating the prevalence of non-zero cells and the average expression of non-zero gene expression[11]. To minimize model bias, the closest 1500 cells per cell type, with the exception of astrocytes, which only presented with 1032 cells[12]. A single cell mixture matrix was constructed of 3000 genes and 6 cellular units and submitted to CIBERSORTx to create a deconvolution model[13].

The resulting deconvolution model was applied to individual DSP ROI data, by generating singular value decomposition (SVD) based knockoff models[14]. These knock off models consisted of 30 psuedo-replicates for each sample which were collated, and deconvolution was performed for each ROI. A Wilcoxon rank sum test was applied the cell type abundances to elucidate any proportional differences.

Differential expression analysis between APP genotypes using the limma R package. Additional sample filtering was applied to the individual cell-type expression models, removing samples presenting with low expression (log2(median) < 4). Outlier samples were identified and removed using the R base boxplot function of the first two principal components. Gene set over representation (ORA) analysis was performed with enrich (v 3.2)[15].

### Visualizations

Information regarding ligand receptor (LR) pairs was obtained from CellTalkDB (http://tcm.zju.edu.cn/celltalkdb/download.php) [16]. LR networks were constructed with igraph (v 2.1.1), and chord diagrams from circlize (v 0.4.16) [17–19].

## Results

GeoMx digital spatial profiling (Bruker NanoString, lnc.) was performed on12 aged (11 month) mouse models. One thirty-three ROIs were collected, 79 from App-wildtype mice (32 ROIs from female and from 39 male) and 62 from App+ transgenic (30 female and 32 male) mice. An example ROI is shown in Figure 1. Both genotype and gender were equally represented among the mouse models and ROIs (Fisher’s test, p-value=0.73). An average of 11.1 ROIs per mouse model was observed. The least represented mouse model presented 6 usable ROIs and the most represented mouse model yielded16 ROIs. One area of illumination AOI was collected from each ROI.

**Figure 1:**
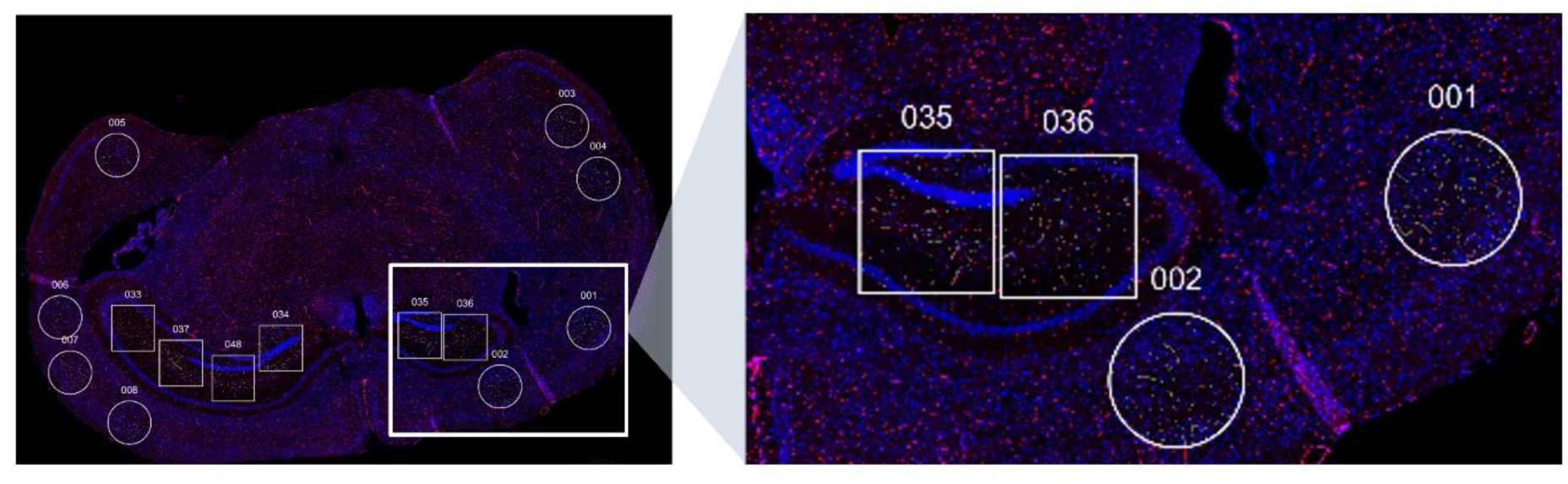
ROI selection. Regions of interest (ROIs) were selected for digital profiling analyses from slides of Alzheimer’s disease transgenic mice and wild-type control mice. Circular ROIs of *660* μm *in diameter* were drawn in the cortex, and square ROIs of 660 µm x 700 μm were drawn in the hippocampus. The slides were imaged for CD31 (red) to visualize endothelial cells and SYTO-83 (blue) to visualize tissue morphology.

### ROI analysis

RNA sequencing was performed on the ROIs for 19,963 target genes. The modified LOQ, see above methods, allowed for the differential expression (DE) analysis of 7,687 target genes. The DE analysis of APP-and APP+ on these genes were performed using a linear regression model incorporating a fixed effect for gender (Figure 2A). A total of 4,273 genes were found to be statistically significant (p-value <0.05). Subsequently, we evaluated the ROI expression profiles of 7,687 target genes with UCell signature scoring to investigate whether there was evidence of an enriched source of endothelial specific expression. While we observed the expression to be enriched in vasculature cells (endothelial, pericytes, and vascular smooth muscle cells), we observed substantial non-endothelial-based expression, particularly from neuronal cells, as shown in Figure 2B. We then evaluated the cell type composition with a gene signature enrichment analysis of the 4,273 differential expressed genes (Figure 2C) and the overlaps of the observed DE genes with a particular cell type signature (Figure 2D).

**Figure 2:**
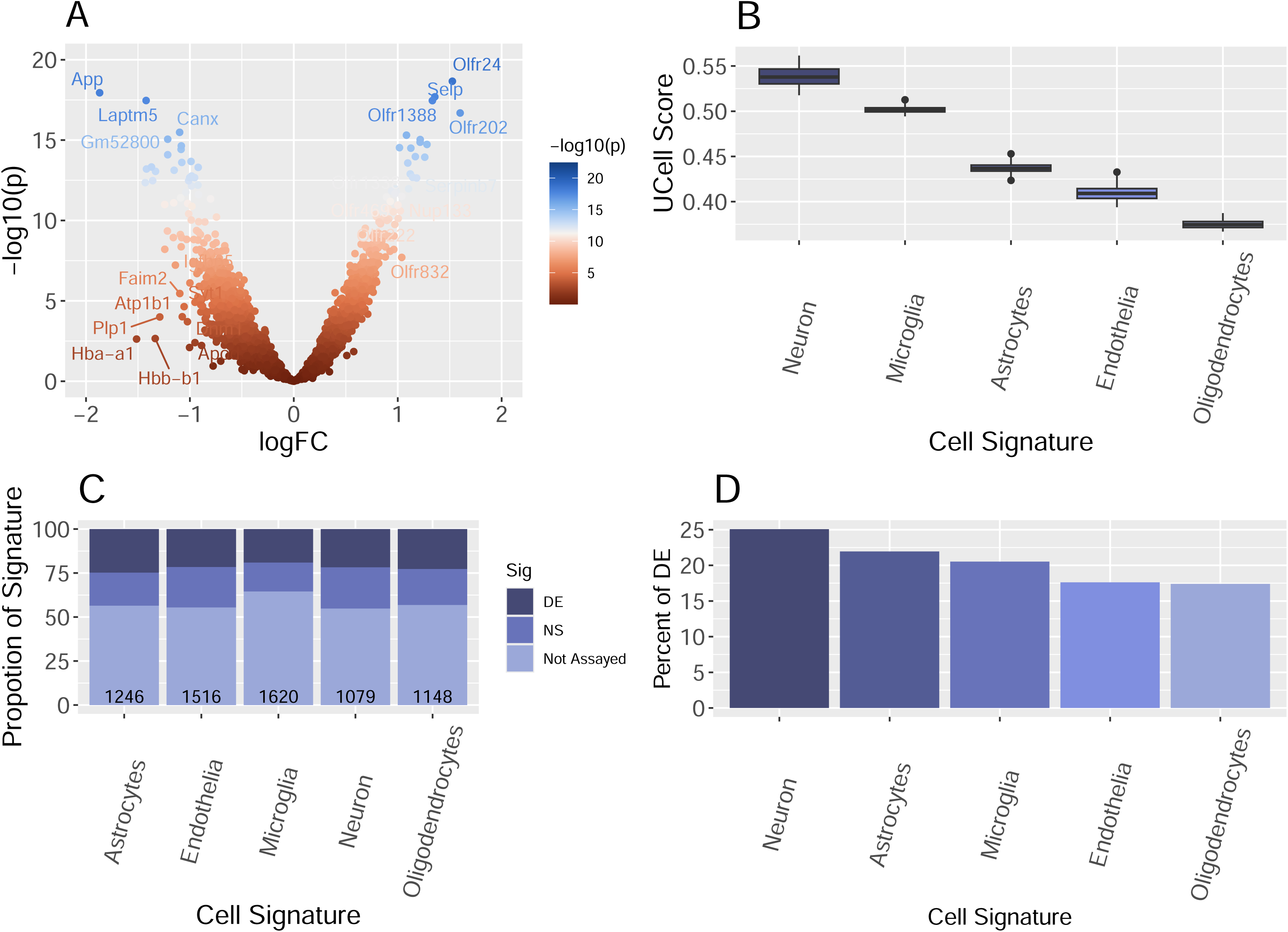
Analysis of collected regions of interest (ROIs). A) A volcano plot illustrating the results of differential expression analyses which identified 66 significant genes, following False Discovery Rate (FDR) correction and each exhibiting a fold change of 2. These significant genes are labeled in the plot. Cell type specific signature analysis was performed on the ROI’s. We observed that although stained for endothelial cells, the ROI expression profiles were influenced by non-endothelial cellular expression. B) Assessment of whether any of the cell type signatures presented with an enrichment of differentially expressed genes. D) We observed that among the differentially expressed genes, there was not a specific cell type source of differential expression.

### Deconvolution Model

Since we did not observe an endothelial-specific signal, we employed a CIBERSORTx deconvolution model developed from 20,333 scRNA cells harvested from 8 aging mice (GSE129788), as described in the Methods section. The deconvolution model, representing 14 cerebrovascular cell types (Figure 3A), was applied to the 133 ROIs generated in our study. In validating the cell types, we observed the co-clustering of endothelial cells with astrocytes and pericytes (Figure 3B) and evaluation of NVU cell type-specific signatures scores with UCell indicated that several cell types exhibited similar scoring patterns. Specifically, we observed the pericytes (PC), vascular smooth muscle cells (VSMC) and endothelial (EC) all scoring the highest with the endothelial signature, and astrocytes (ASC) and astrocyte-restricted precursor cells (ARP) scored for the astrocyte gene signature. Similarly, neuroendocrine cells (NendC) and mature neurons (mNEUR) scored for the neuron gene signature and oligodendrocytes (OLG) and oligodendrocyte precursor cells (OPC) scored high for the oligodendrocyte’s gene signature, suggesting that an overfitting of the original clustering (Figure 3C). Principal component analysis of the cell type signature scores derived from UCell supported the consolidation of these cells into unified cellular units. Additionally, oligodendrocyte precursor cells were found to cluster distinctly from oligodendrocytes and were retained as a separate cell type (Figure 3D). CIBSERSORTx was then utilized to construct a new deconvolution model of the 6 consolidated NVU cell types (Figure 3E).

**Figure 3:**
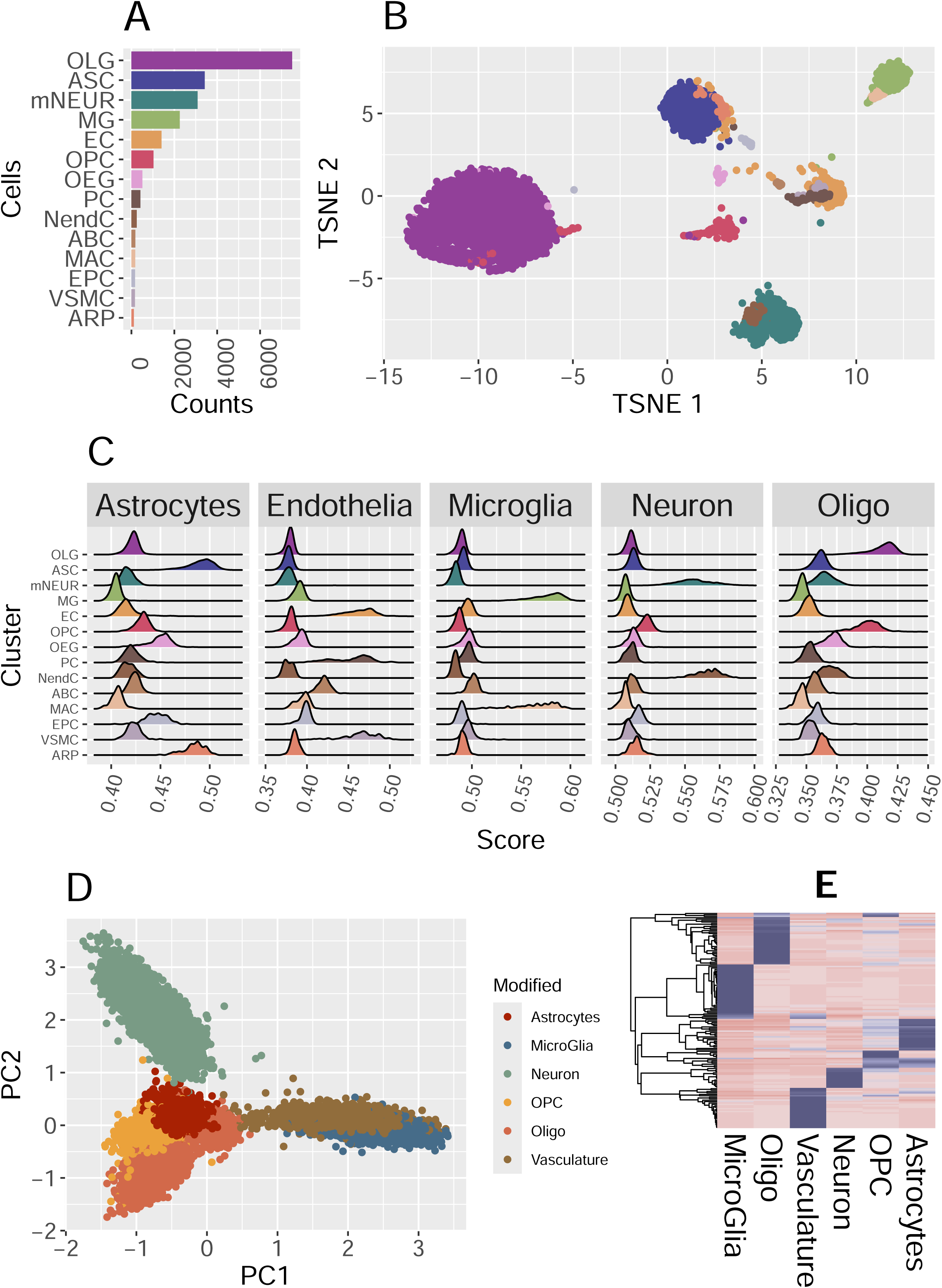
Deconvolution Model (top to bottom). 3A) Single c ell data of 21–22-month-old mice obtained from GSE129788. Fourteen cell types were reported with a predominance of oligodendrocytes. 3B) Visualization of cell type clusters demonstrated co-clustering. 3C) UCell scoring of NVU cell type signatures observed several cell types demonstrating similar and correlated scoring patterns. The pericyte, endothelial, and VSMC single cells all scored high for the endothelial signature. 3D) PCA of UCell scoring matrix indicating the collapsing of 14 cell types into 6 cellular units, and the removal of under-represented EPC and ABC cells. 3E) CIBERSORTx generated deconvolution model.

### ROI Cell type prevalence

Among the 6 cellular units we collected within the ROIs we observed a significant increase in the prevalence of microglial cells among the APP+ ROI’s (p-value 8.6X10^^-3^), Figure 4A. More importantly, we observe that the endothelial-enriched vasculature is the most prevalent cell type represented in the ROI selection, with a median proportion of 31.6%. Neurons and astrocytes were the next most prevalent cell types, with medians of 28.0% and 25.0%, respectively. The neuron and endothelial prevalence demonstrated the strongest correlations (Spearman’s-0.81), exceeding that of the OPC-astrocytes (-0.49) and neuron-astrocytes (-0.42). The OPC, oligodendrocytes, and microglia all presented with median proportions of less than 5%, as shown in Figure 4A.

**Figure 4:**
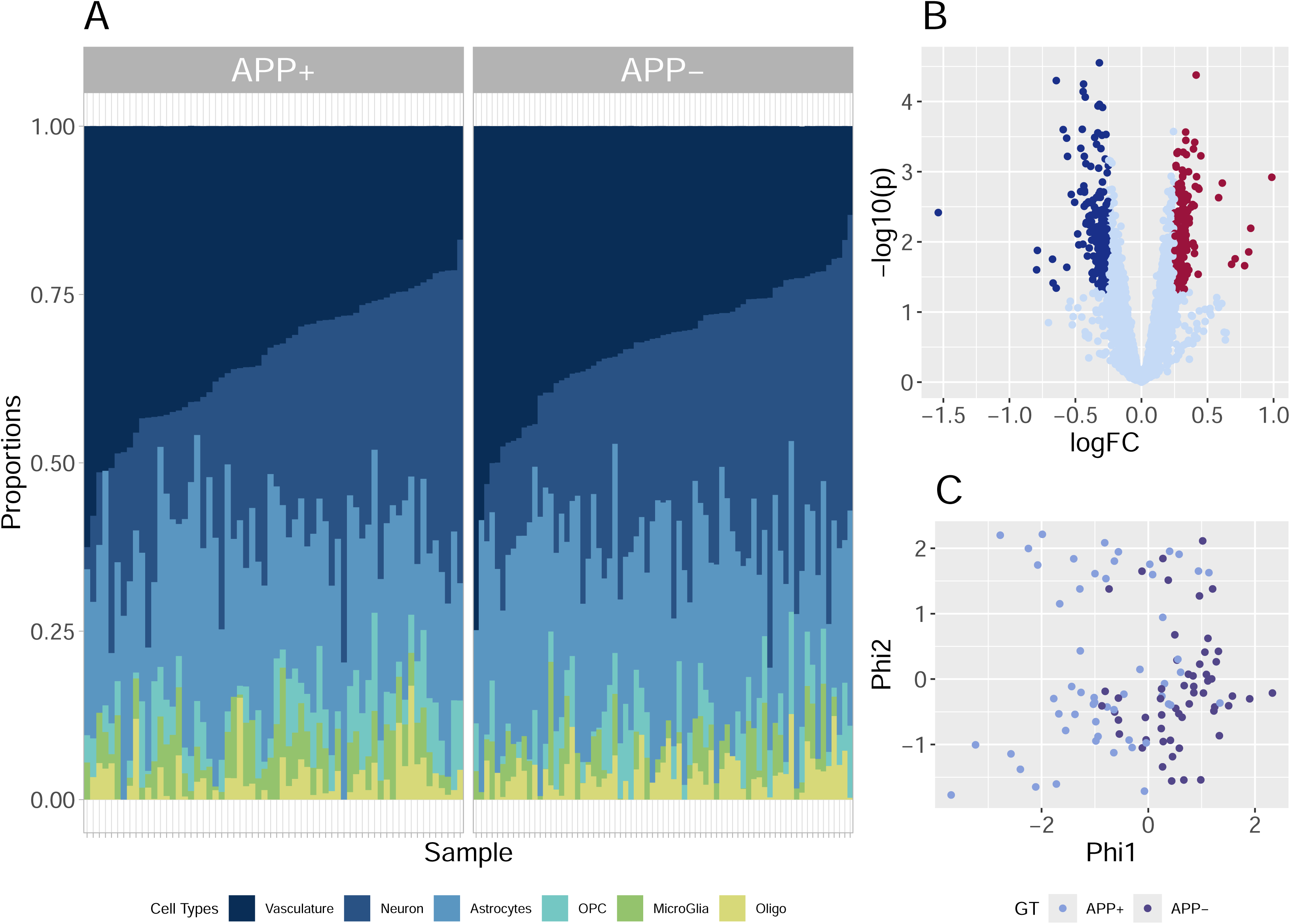
Analysis of deconvolved ROIs. A) The proportions of the 6 cell types observed in the133 ROIs are presented in these stacked bar charts. The cell types were ordered by their median proportions and samples are ordered by the median proportion of endothelial-enriched vasculature. The strong negative correlation (-0.81) between neuronal and endothelial prevalence is notable in the diagram. B) A volcano plot of the differential expression observed in the endothelial-enriched vasculature. Differential expressed genes (p < 0.05) presenting with a logFC > 0.25 are indicated in red (247 genes) and Differential expressed genes (p < 0.05) presenting with a logFC > 0.25 are indicated in blue (213 genes). C) Lafon’s dimension reduction using the 460 DE genes, demonstrating the separation of APP+ samples from APP-samples.

### Cell type differential expression

To gain insight into the transcriptional differences of individual NVU cell types that underlie the APP genotypes, we performed cell-type specific DE analysis on the deconvolved transcription data for 120 (of 133, 90.2%) of the ROIs by implementing SVD knock-off models of the individual ROIs. Thirteen ROIs were excluded from the analysis due to instability observed in the deconvolution model when 2 or more NVU cell types were missing in these samples. Differential expression was assessed between the APP genotypes adjusting for gender, Supplementary File 1. Expression variability is notably reduced in the deconvolved expression profiles. A volcano plot of the differential expression in the endothelial-enriched vasculature was shown in Figure 4B. We observed 993 DE genes (p-value< 0.05) in the endothelial-enriched vasculature, 460 (247 up-regulated in red and 213 down-regulated in blue) presented with |logFC| > 0.25. Figure 4C presents a diffusion-based Lafon’s dimension reduction using an SVD of Spearman’s correlation of the 460 DE genes. The imputed cell type data presents with less variance than assayed data, and Lafon’s method better captured the latent structure among the samples over a standard PCA. The samples demonstrate genotype-specific clustering along the first Phi dimension.

Gene set ORA was performed on the 993 differentially expressed genes against 271 KEGG gene sets. Fourteen gene sets were considered significantly enriched after adjusting for multiple testing, provided in Figure 5. Among these gene sets, the most significant gene sets were associated with maintaining electrolyte homeostasis (Supplementary File 2). Additional gene sets observed were associated with cellular signaling.

**Figure 5:**
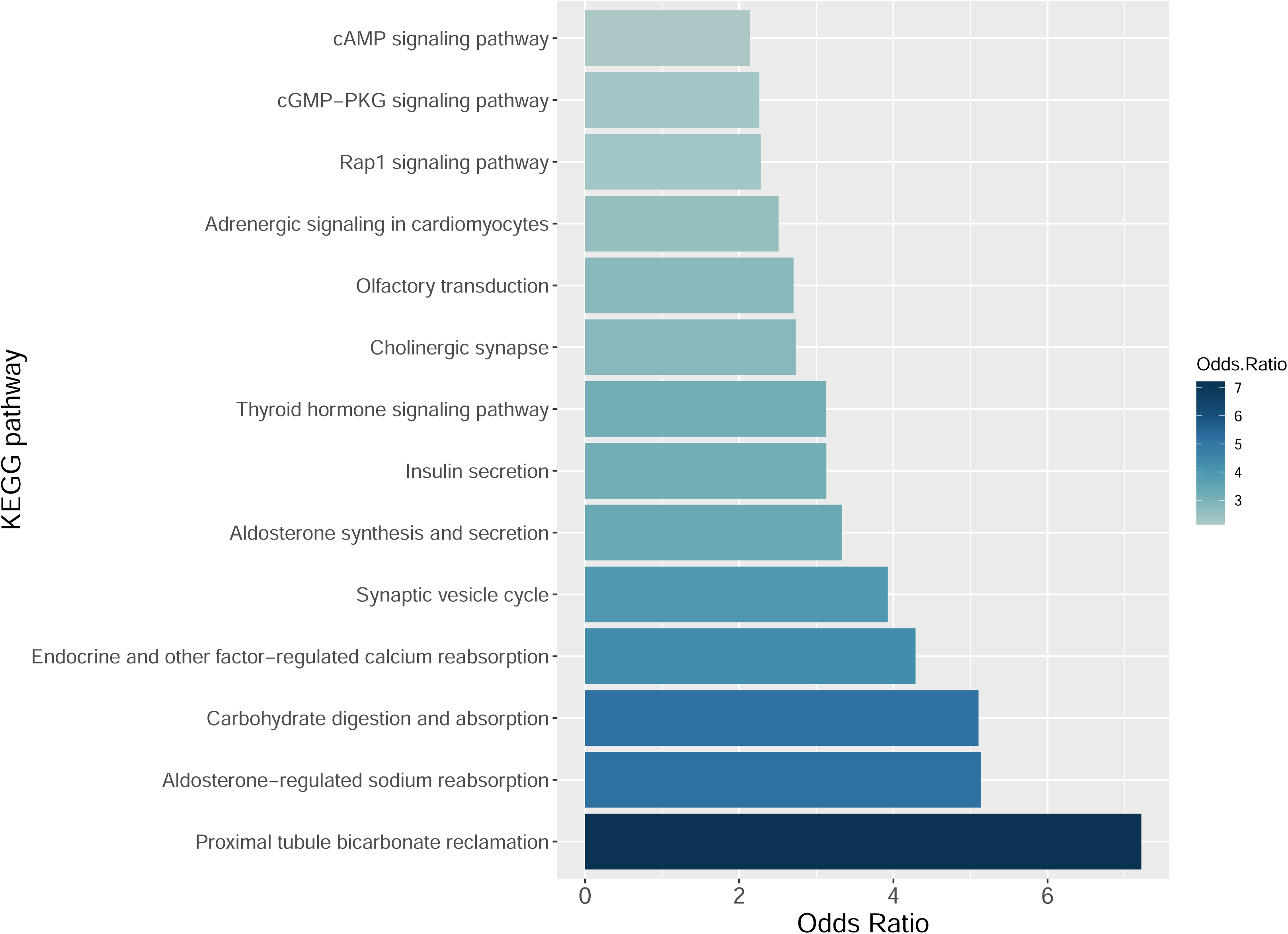
KEGG gene set enrichment. Gene set ORA enrichment was performed using the 993 differentially expressed genes from the endothelial-enriched vasculature. The 993 genes were associated with 271 KEGG gene sets, 47of which were observed to be significant (p-value <0.05). FDR p-value correction was applied, resulting in the 14 gene sets depicted in the Figure 5 barplot. The 14 gene sets are presented in increasing order of the Odds Ratio associated with their over representation. Median Odds ratio was 3.1 (95% confidence of 2.7 to 4.4).

### Ligand-receptor (LR) networks of the NVU

Given the observed over-representation of DE genes associated with cellular signaling we investigated potential LR interactions which were evident in the cell specific data.

Using sample-specific imputed cell types, we identified known LR pairings that were both significantly up-regulated and with a Spearman’s correlation greater than 0.35. As expected, fewer pairings were observed with oligodendrocytes (Oligo) and precursor cells (OPCs), which are not components of the NVU (Supplementary File 3). The observed NVU LR pairings were tallied and presented in Figure 6A. Not surprisingly, given the regulatory function of maintaining the BBB integrity, we observed prominent astrocyte mediated communication among the NVU components. The endothelial-enriched vasculature provided the second most common source of mediated signaling events, suggesting a feedback role for the vasculature.

**Figure 6:**
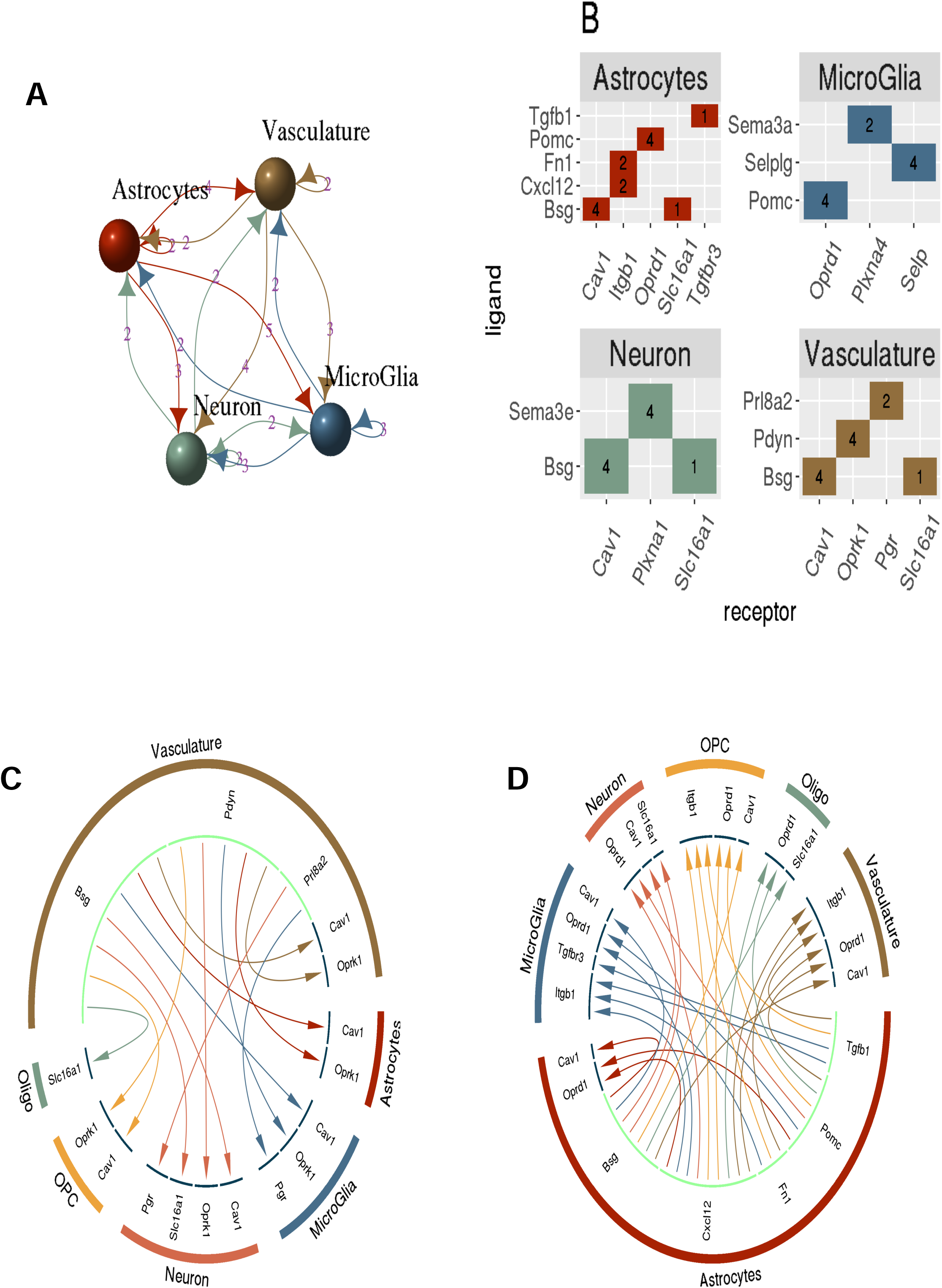
The specific ligand-receptor relationships observed among the NVU. A) A graph depicting the cell types of the NVU, and the number of ligand-receptor pairings. B) The observed ligand-receptor interactions among the deconvolved cell types. For the endothelial-enriched vasculature we observed the Oprk1 ligand was observed to interact with the Pdyn receptor for all 4 cell types, including the autocrine signaling with itself. C) A chord diagram depicting the potential feedback of the vasculature-enrich ligand signaling effects in response to astrocyte regulation. D) The chord diagram indicating the regulatory signals originating from the astrocyte

The ligand-receptor pairs identified for each NVU cell type are detailed in Figure 6B. We tallied all observed pairs among the three primary NVU cell types respectively and included the autocrine signaling pair of the whole vasculature unit. Chord diagrams were constructed to present the ligand-receptor pairings from the regulatory astrocytes and the feedback from the vasculature. The potential feedback mechanism could be observed via the astrocyte mediated Basigin (Bsg) ligand signaling which was observed to pair ubiquitously across the NVU with Cav1 and the vasculature providing the same ligand-paired expression in return to the NVU cell types (Supplementary File 4).

Similarly, the endothelial-enriched vasculature Prodynorphin (PDYN) ligand paired with the kappa opioid receptor (Oprk1) receptor in NVU cell types, see Figure 6C. The same regulatory loop was not observed coming from the astrocytes, rather a ligand pairing of Pomc with the NVU bound delta opioid receptor (Oprd1), see Figure 6D.

## Discussion

We designed our experiment to utilize digital spatial profiling of the endothelial component of the vasculature that comprises the blood-brain barrier. The assessment of the expression profiles of the ROIs indicated that significant contribution from non-endothelial cells (Figure 2B-2D). To address this, we constructed and applied a deconvolution model (Figure 3A–3E) to provide a focused assessment of the endothelial-enriched vasculature (Figure 4A).

Bioinformatics analysis of the dysregulation in the APP+ endothelial-enriched vasculature revealed genes associated with cellular signaling, as well as carbohydrate digestion and absorption pathways. The upregulation of Pi3kcd, Pik3cb, and Akt1 genes corroborates with the existing literature that Pi3k/Akt signaling is prevalent in patients with dementia[20]. Similarly, upregulation was observed in Map2k3 and Tln1, members of the Rap1 signaling pathway which is crucial for cell adhesion and regulates inflammation and lymphocytic migration Specifically, Tln1 a VLA-4 affinity regulator, has been implicated in both spontaneous and chemokine-triggered rapid adhesions to VCAM-1[21] which has been associated with cerebrovascular inflammation[22] and the transmigration of leukocytes[23, 24].

Several other signaling pathways were observed to be disrupted in APP+ mice, (Figure 5), including calcium homeostasis and insulin signaling in the brain [25, 26].

Specifically dysregulation was observed in Pik3cg, Ins, and Akt1, which are involved in regulating brain insulin signaling[27, 28]. While upregulation of Camk2b, Ppp3cb, and Plcb1 suggest that calcium signaling was upregulated in the APP+ mice. Camk2b has been reported to be involved in regulating synaptic plasticity[29, 30], Aβ production[31]. Finally, dysregulation of Atp1a3, Atp1a2, Atp1a1, and Atp1b1, suggests disruptions in ion homeostasis and neuronal function in the APP+ model.

Moreover, we investigated the paracrine and autocrine ligand-receptor signaling within the NVU. This identified a feedback loop of astrocyte mediated signaling through of the transmembrane glycoprotein, CD147 (Bsg), and the caveolin-1 (Cav-1) receptor. Both molecules have been implicated in the regulation of Aβ [32, 33] [34]. This signal loop was reciprocated via the endothelial-enriched vasculature, Figure 6C and 6D. A potential second regulatory loop was observed with through the astrocyte Pomc ligand and NVU delta opioid receptors (Oprd1) receptors. Conversely, the vasculature feedback instantiates with the Prodynorphin (Pdyn) ligand and involved the sister kappa opioid receptor (Oprk1) species in the cells of the NVU. Dynorphins have been reported to have elevated levels in postmortem samples from AD patients with increases in dynorphin levels linked to learning and memory impairments and cognitive defects [35–37] The feedback communication observed among the cells in the NVU need to be investigated further, as the offer new therapeutic targets for treatment of the disease.

## Footnotes

This work was supported by the Minnesota Partnership for Biotechnology and Medical Genomics [Grant 00056030], National Institute of Health

[Grant AG058081], and National Institutes of Health/National Institute of Neurological Disorders and Stroke [R01NS125437].

## Acknowledgements

Funding was provided by National Institute on Aging grant [R01AG085900 to K.R.K., X.T. and K.J.T.]

Graphical abstract was created on Biorender.com.

## Data Availability Statement Samples

The authors declare that all the data supporting the findings of this study are contained within the paper.

## Authorship contributions

*Participated in research design:* V.S.S., K.K.K, K.R.K., K.J.T, X.T., and V.J.L.

*Conducted experiments*: V.S.S. and G.L.C.

*Contributed to new reagents or analytic tools:* K.K.K, V.J.L., and K.R.K.

*Performed data analysis*: V.S.S. and K.J.T., *Wrote or contributed to the writing of the manuscript:* V.S.S., K.J.T., K.R.K, and K.K.K.

