## Supplementary figures and images for "Characterization of transcriptomic changes in the neurovascular unit of Alzheimiers transgenic mouse models using digital spatial profiling"

### Supplementary File 3

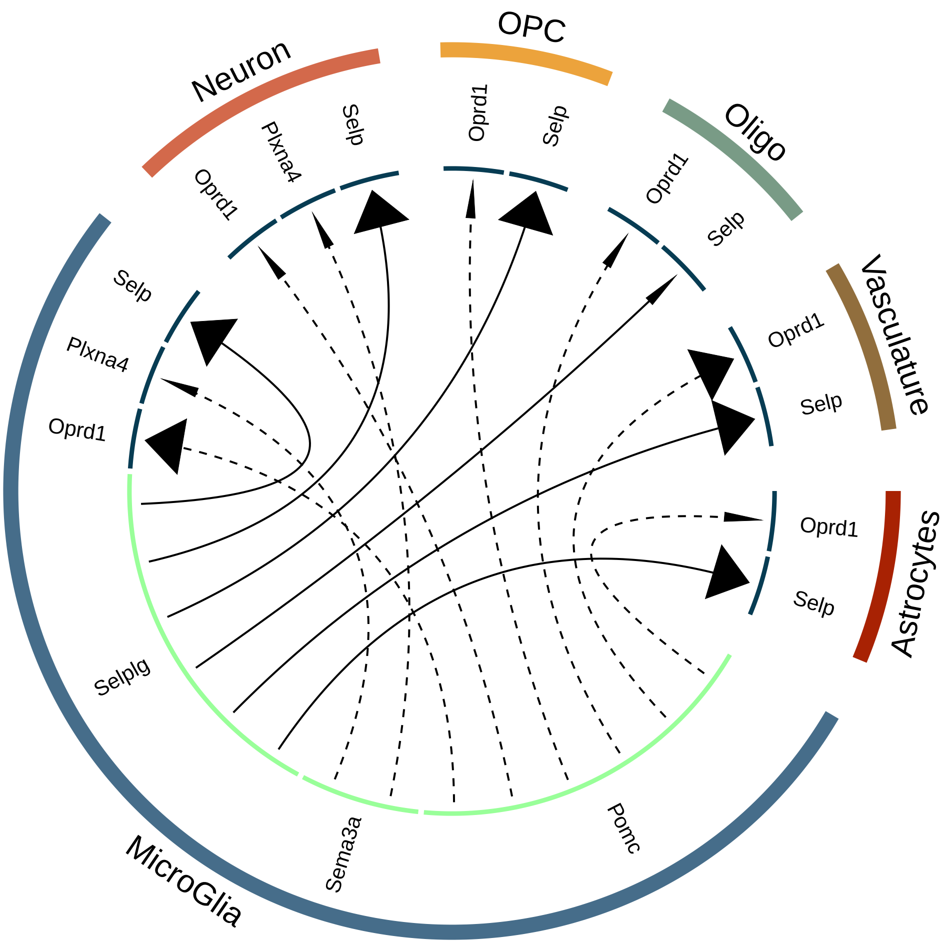

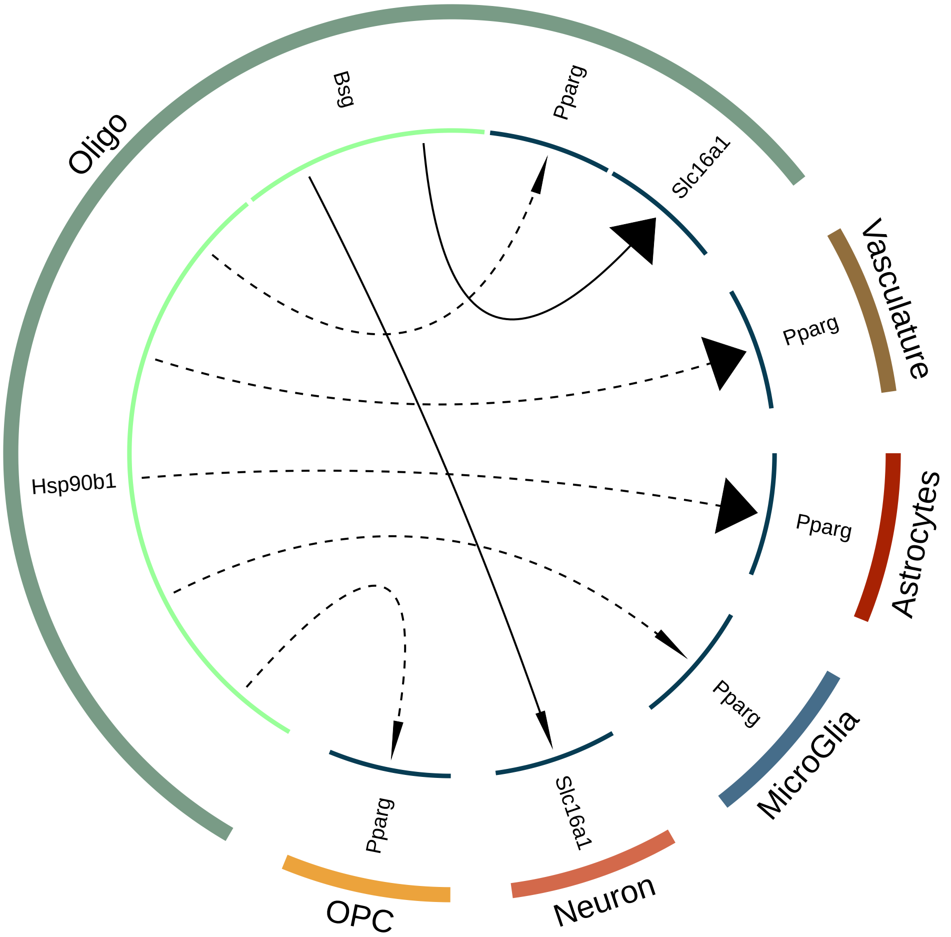

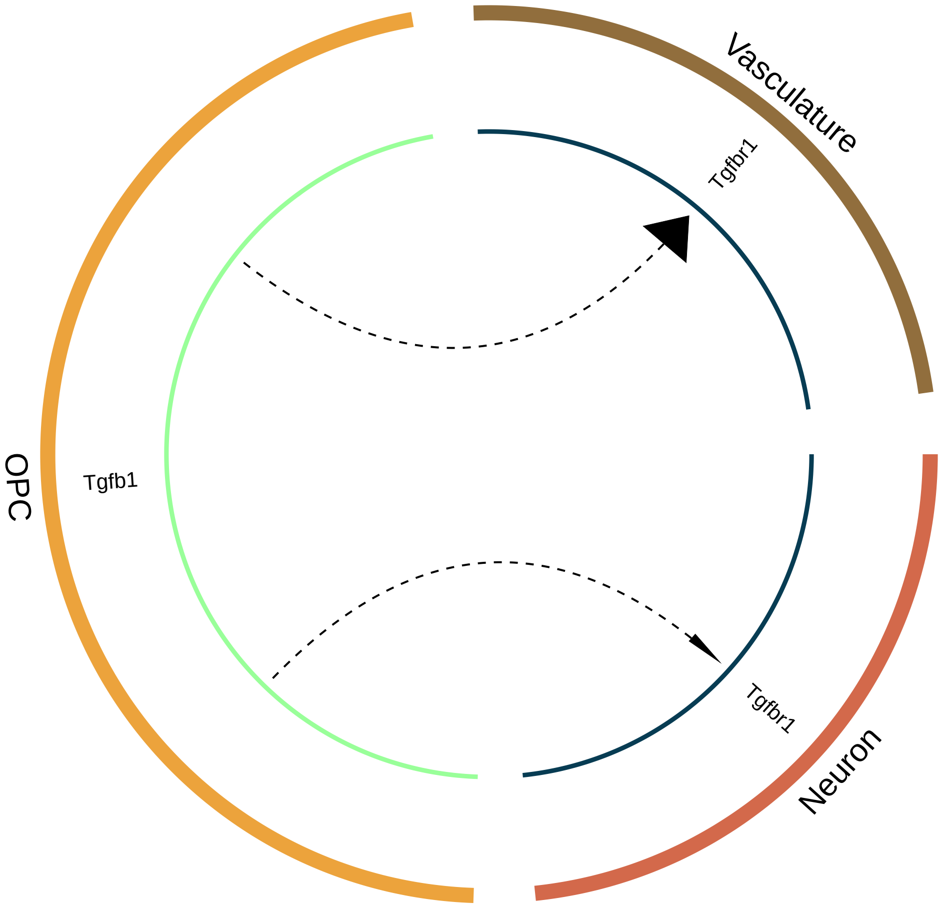

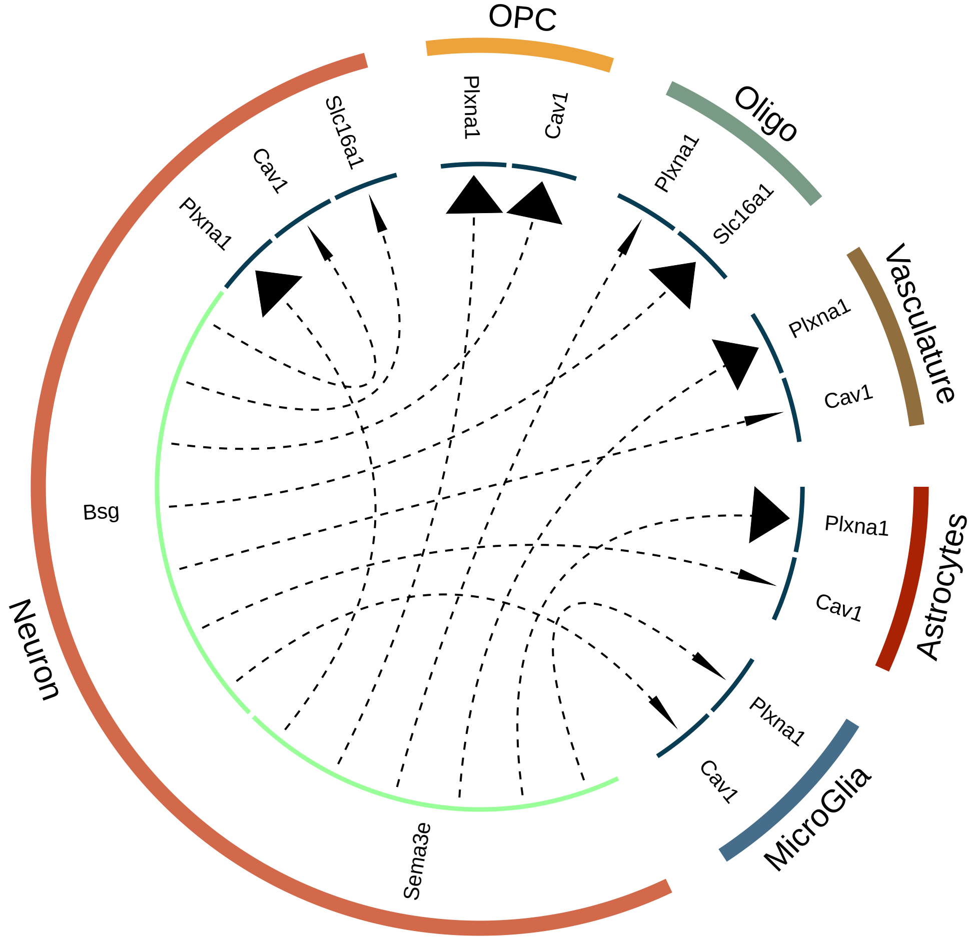

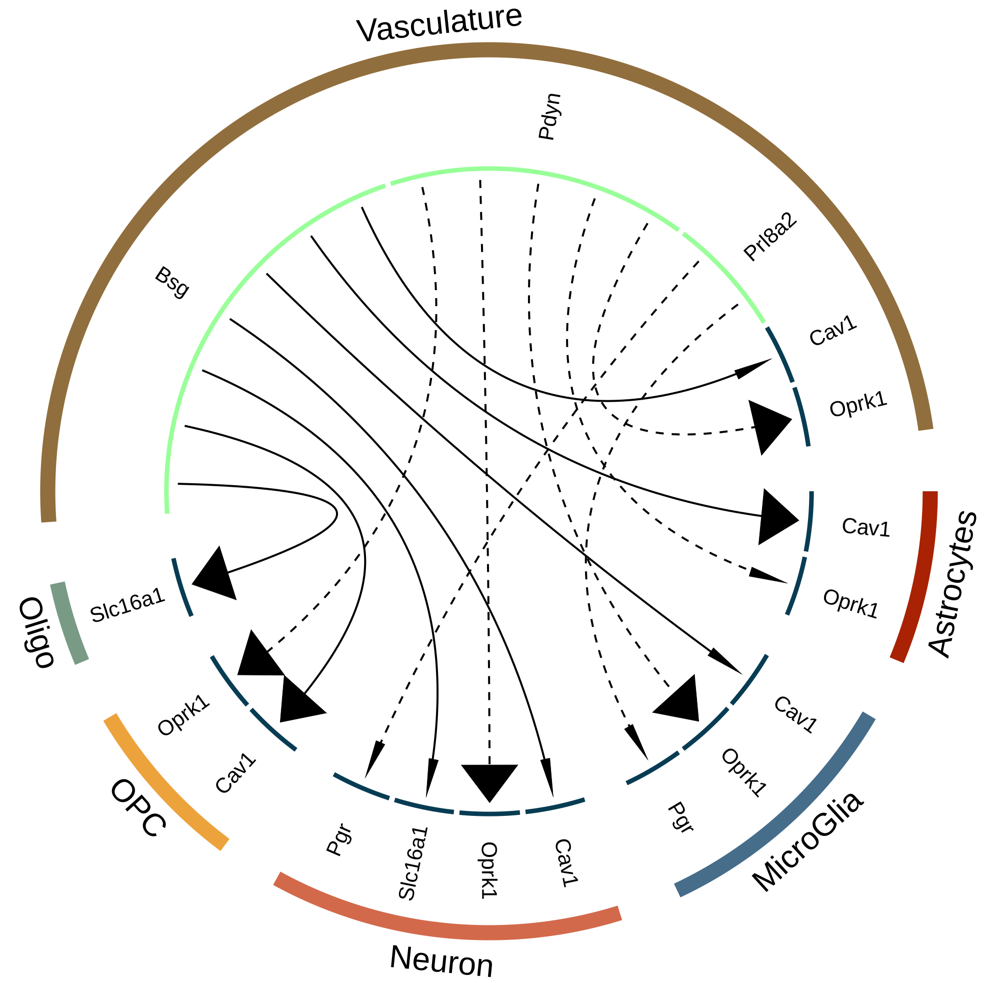

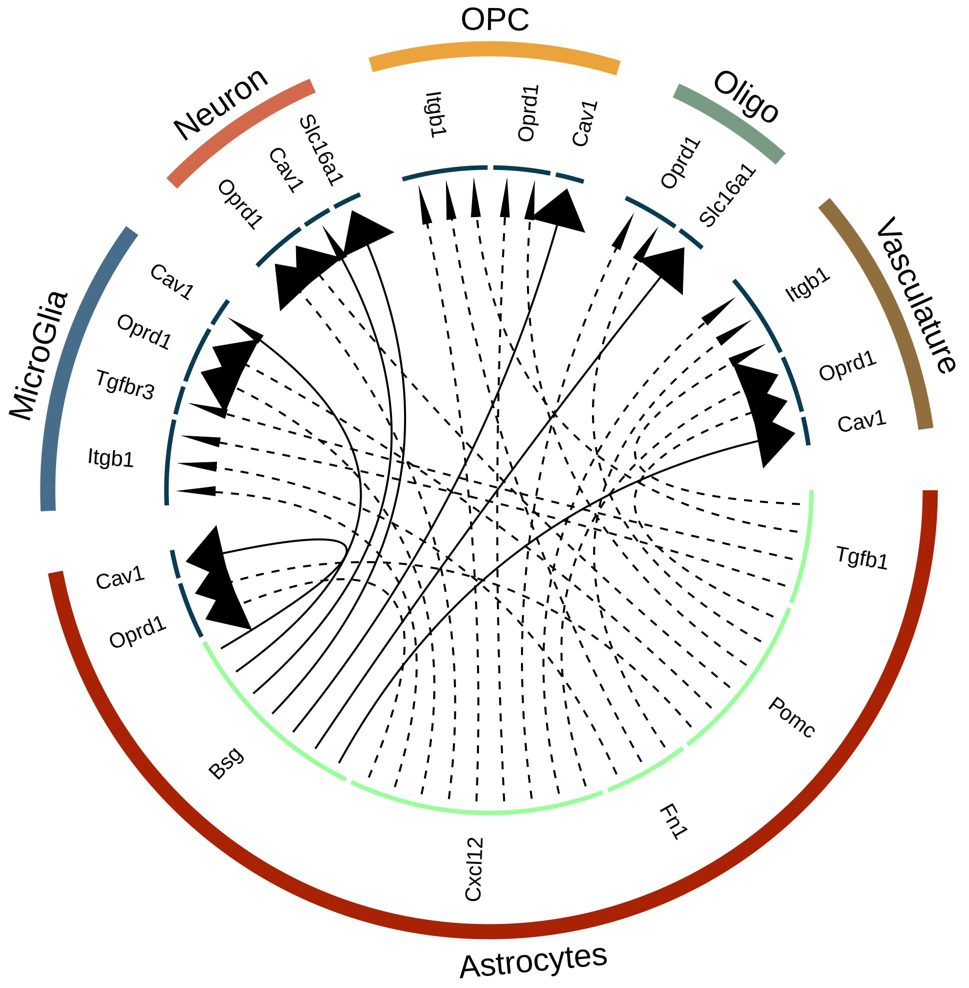
